# Deep Generative Design with 3D Pharmacophoric Constraints

**DOI:** 10.1101/2021.04.27.441676

**Authors:** Fergus Imrie, Thomas E. Hadfield, Anthony R. Bradley, Charlotte M. Deane

## Abstract

Generative models have increasingly been proposed as a solution to the molecular design problem. However, it has proved challenging to control the design process or incorporate prior knowledge, limiting their practical use in drug discovery. In particular, generative methods have made limited use of three-dimensional (3D) structural information even though this is critical to binding. This work describes a method to incorporate such information and demonstrates the benefit of doing so. We combine an existing graph-based deep generative model, DeLinker, with a convolutional neural network to utilise physically-meaningful 3D representations of molecules and target pharmacophores. We apply our model, DEVELOP, to both linker and R-group design, demonstrating its suitability for both hit-to-lead and lead optimisation. The 3D pharmacophoric information results in improved generation and allows greater control of the design process. In multiple large-scale evaluations, we show that including 3D pharmacophoric constraints results in substantial improvements in the quality of generated molecules. On a challenging test set derived from PDBbind, our model improves the proportion of generated molecules with high 3D similarity to the original molecule by over 300%. In addition, DEVELOP recovers 10 × more of the original molecules compared to the base-line DeLinker method. Our approach is general-purpose, readily modifiable to alternate 3D representations, and can be incorporated into other generative frameworks. Code is available at https://github.com/oxpig/DEVELOP.

## Introduction

Drug design optimises molecules through a multistep, iterative process in order to achieve a desired biological response. The size of the search space^1^ and discontinuous nature of the optimisation landscape^2^ are two key factors contributing to the difficulty of this problem and, as a result, currently molecular design is typically led by human experts.

Machine learning models for molecule generation^3–5^ offer an alternative approach to human-led design or rules-based transformations.^6,7^ Despite recent success,^8^ for these methods to be broadly adopted in drug discovery, more control over the generative process is required, including the ability to incorporate prior knowledge. In the hit-to-lead (or lead generation) and lead optimisation stages of drug discovery, the goal is to improve one, or several, properties. This is typically achieved by modifying an existing molecule rather than designing a compound from scratch. Such modifications can be broadly categorised into one of two scenarios: linker design and scaffold elaboration.

Linker design is a general problem in drug discovery capturing a wide range of tasks where the goal is to design a molecular scaffold that incorporates two (or more) specific substructures. Three key applications that can all be considered as linker design are scaffold hopping,^9,10^ fragment linking, ^11,12^ and PROTAC design.^13,14^ Examples of these design tasks are shown in Figure 1a–c.

**Figure 1:**
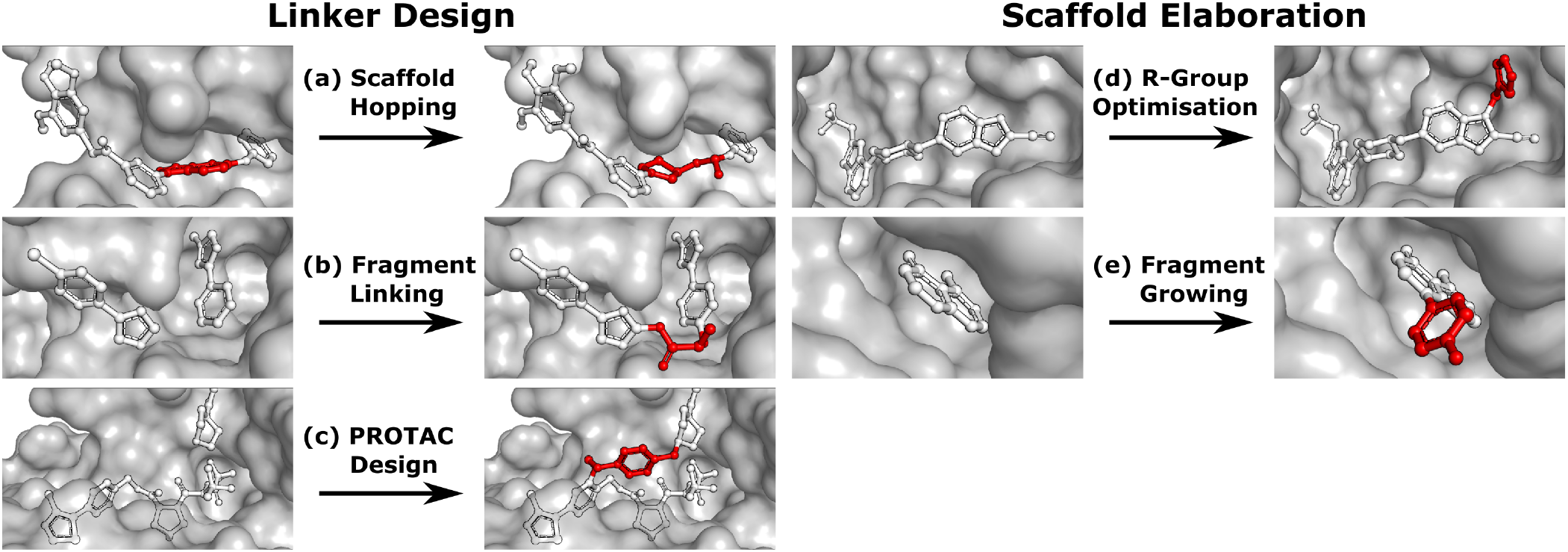
Design tasks considered in this work. (a)-(c) cover linker design, specifically (a) scaffold hopping, (b) fragment linking, and (c) PROTAC design. (d)-(e) scaffold elaboration, namely (d) R-group optimisation and (e) fragment growing. Components of the ligand that are modified or added in the design process are shown in red.

In contrast to linker design which is tasked with discovering molecular cores, scaffold elaboration proposes molecules incorporating these privileged substructures. Scaffold elaboration covers a broad range of medicinally important scenarios, such as R-group optimisation ^15^ and fragment growing^12,16^ (Figure 1d–e). R-group optimisation is utilised in almost all drug discovery projects to improve the potency, selectivity, and other properties of a molecule during lead optimisation and characterise the structure-activity relationship (SAR) of a molecular series, while growing is the primary method for elaborating fragment hits.

Recently, several deep learning methods have been proposed to address these design challenges. We previously published the first application of deep learning for molecular linker design (“DeLinker”), ^17^ reporting substantial improvement over a database-based approach, the previous *de facto* computational method for this task, by including basic structural information. Yang et al. ^18^ have since proposed an alternative model (“SyntaLinker”) based on the transformer architecture and a SMILES-based representation. Their model did not incorporate structural information but instead included 1D molecular patterns capturing factors such as the shortest linker bond distance.

Deep learning approaches have also been proposed for scaffold elaboration. Graph-based approaches were proposed by Lim et al. ^19^ and Li et al. ^20^. The scaffolds employed in both methods do not have explicit attachment points. As such, these methods are primarily applicable to the general generation of molecules with a privileged scaffold or substructure, rather than tasks such as R-group design. In contrast, Arús-Pous et al. ^21^ developed a preprocessing formulation to permit a SMILES-based approach that requires specific attachment points to be defined. In both linker design and scaffold elaboration, some knowledge about the desired modification is typically available;^22^ this can either be derived from the protein binding site in the case of structure-based design,^23^ or from other molecules in ligand-based drug discovery.^24^ In either case, this information has strong 3D dependencies which should be taken into account. However, currently this information, which is crucial to successful compound design, is typically not utilised by generative models.^25^

None of the existing machine learning models for linker design or scaffold elaboration effectively utilise structural information, with DeLinker the only framework explicitly incorporating any 3D information in the form of the distance between the starting substructures and their relative orientations. While this minimal parametrization had a substantial impact on the quality of the generated molecules,^17^ much of the key information about the characteristics of the binding site is not taken into account in the generative process.

There have been several recent approaches proposed to generate molecules from 3D representations. Skalic et al. ^26^ generated molecules from a 3D representation of a seed ligand. However, their approach requires a known active molecule, only provides 3D information implicitly to seed their model, and offers no further control over generated compounds. As a result, their generative model recovered fewer than 2% of seed molecules. This idea was extended in Skalic et al. ^27^ to generate the ligand representation from the protein target. While this alleviates the need for a known active, it is not possible to use prior knowledge to influence the ligand representation. Finally, in concurrent work to this paper, both Ragoza et al. ^28^ and Masuda et al. ^29^ generate molecules by adopting an autoencoder framework to first generate atomic densities, before using a fitting procedure to convert the continuous 3D grids to discrete molecular structures.

All prior approaches utilising 3D representations attempt to generate entire molecules and do not readily incorporate expert knowledge. While this is arguably the end-goal for molecular design, in practice this limits the applicability of such methods. In particular, it prevents their use in later stage drug discovery where there is significant prior knowledge that could and should inform compound design.

In this paper, we propose DEVELOP (DEep Vision-Enhanced Lead OPtimisation), a graph-based generative model that uses a convolutional neural network (CNN) to incorporate physically-meaningful 3D structural information, here provided as 3D pharmacophores,^30^ a general and widely-used representation in cheminformatics. Our model is applicable to a wide variety of design tasks in the hit-to-lead and lead optimisation stages of drug discovery, covering linker design and scaffold elaboration. Importantly, the richer representation of the binding site readily and naturally allows the incorporation of domain knowledge and significantly improves the quality of generated compounds. On a challenging test set derived from PDBbind, our model improves the proportion of generated molecules with high 3D similarity to the original molecule by over 300%. In addition, DEVELOP recovers 10 × more of the original molecules compared to the baseline DeLinker method.

## Methods

This work describes DEVELOP, a deep learning approach combining GNNs and CNNs for molecular linker design and scaffold elaboration. We extend current molecular generative methods to incorporate physically-meaningful 3D structural information, enabling prior knowledge to be readily incorporated and allowing greater control of the generative process by domain experts. Our underlying model is based on Imrie et al. ^17^, which built on the generative process introduced by Liu et al. ^31^ that constructs molecules “bond-by-bond” in a breadth-first manner. Here we outline the generative process and describe how 3D structural information is incorporated (Figure 2).

**Figure 2:**
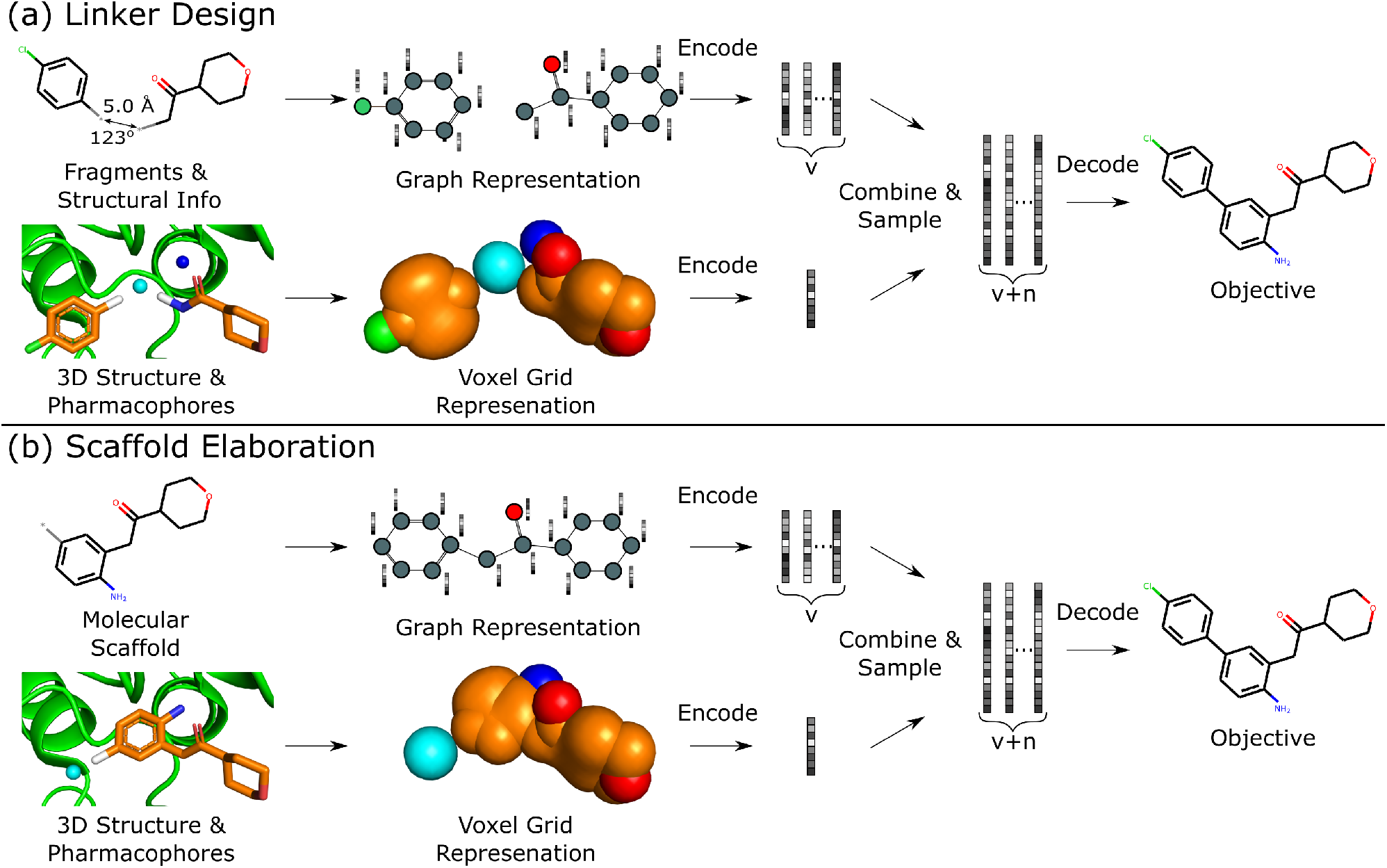
Overview of DEVELOP. The starting structures and 3D pharmacophore map are converted into a graph representation and a voxel grid, respectively. These are fed into GNN and CNN encoders, respectively. The featurisations are combined and decoded by a GNN-based decoder.

### Generative Process

To perform the generative tasks considered in this manuscript, DEVELOP takes as input (i) the chemical structure of either the substructures that are to be linked or the molecular scaffold that is to be elaborated and (ii) a 3D structure of the partial molecule and the desired pharmacophoric features. The input to DEVELOP can be seen in Figure 2 for both linker design and scaffold elaboration.

Pharmacophores are a widely-used representation in cheminformatics.^30^ They are designed to capture the key chemical interactions that allow ligands to bind to macromolecular targets, such as hydrogen bonds, charges, or lipophilic contacts. Pharmacophores can be derived both from other molecules (ligand-based) and inferred or proposed based on the protein target of interest (structure-based), making this representation broadly applicable. In this work, we utilised 3D pharmacophores derived from the ground truth molecules.

Due to their prevalence and importance in drug discovery, the pharmacophores included in our representation were hydrogen bond donors, hydrogen bond acceptors, and aromatic systems. Our framework is readily extendable to additional pharmacophores, or alternate structural representations.

To generate new molecules, first, a graph representation of the starting substructure(s) is constructed and nodes are encoded using a gated graph neural network (GGNN)^32^ in line with Imrie et al. ^17^. The 3D structure of the starting molecular fragment(s) and desired pharmacophores is voxelised to construct a 3D grid, with atoms and pharmacophores adopting a Gaussian representation centered at their input coordinates^33^ (Figure 2). The voxel grid representation is passed into a 3D convolutional neural network composed of three 3 × 3 × 3 convolutional layers with ReLU activation, each followed by a 2 × 2 × 2 max pooling layer, with the final convolutional layer followed by a global max pooling operation. We then apply dropout with probability 0.2 before a fully-connected layer produces the 3D structural encoding.

For linker design, the distance and angle between the starting substructures has been shown to provide a useful constraint.^17^ However, this representation is not readily extendable to scaffold elaboration, and thus this information is only provided for linker design. The 3D structural encoding is concatenated with the distance and angle information (in the case of linker design) and a 1D count vector representing the number of each pharmacophoric feature that should be present in the generated molecule. This forms the structural information, ***D***, used by the decoder to generate molecules.

From these embeddings, molecules are generated in line with Imrie et al. ^17^. The decoding process is initialised with the node encodings together with a set of expansion nodes whose feature vectors are drawn from the standard normal,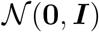. Each node is labeled with an atom type sampled from a classifier applied to the concatenation of the node encoding and the structural information, ***D***.

Molecules are constructed iteratively “bond-by-bond” from this set of nodes. After each step, the node encodings are updated by a decoder GGNN. Edges and their edge types are chosen based on the feature vector for the (possible) edge between node *υ* and candidate node *u* given by

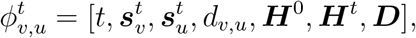

where ***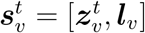*** is the concatenation of the hidden state of node *υ* after *t* steps 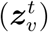 and its atomic label (***l****_υ_*), *d_v,u_* is the graph distance between *v* and *u*, ***H***^0^ is the average initial representation of all nodes, ***H****^t^* is the average representation of nodes at generation step *t*, and ***D*** represents the structural information.

Our model is trained using the same loss function as Imrie et al. ^17^ which is similar to the standard VAE loss, including a reconstruction loss and a Kullback-Leibler (KL) regularisation term:

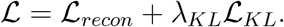

No extra terms are included to regularise the CNN encoding. We use the same hyperparameters for training as Imrie et al. ^17^ (see Supporting Information). For additional details regarding the underlying model see Imrie et al. ^17^.

### Data Sets

Due to the lack of experimental data, we constructed sets for training and evaluation from general molecular data sets using standard transformations from matched-molecular pair analysis.^34^ For both linker design and scaffold elaboration, we used the same underlying data and adopted the same process for constructing datasets, with the main difference the transformation used. For linker design, we enumerated all double cuts of acyclic single bonds that were not within functional groups, while for scaffold elaboration we performed single cuts.

The training sets were derived from the subset of ZINC^35^ selected at random by Gómez-Bombarelli et al. ^3^ using the fragmentation procedure described above. For linker design, this results in c. 418,000 fragment-molecule pairs and is the same training set as Imrie et al. ^17^, while for scaffold elaboration there are c. 427,000 examples.

To evaluate our method, we constructed test sets for linker design and scaffold elaboration from CASF-2016^36^ and the PDBbind Refined Set^37^ (v. 2019) using the same fragmentation procedure used to construct the training set. For both of the CASF and PDBbind test sets, we only retained examples with elaborations containing at least five atoms. In addition, for the PDBbind test sets, we ensured that the molecular elaboration was unique and was not present in the training set. As a result, the CASF test sets contain 188 and 237 examples for linker design and scaffold elaboration, respectively, while the PDBbind test sets contain 321 and 295 examples, respectively. Due to the stricter inclusion criteria, the PDBbind test sets represent a significantly more challenging test than the CASF sets and should better capture the ability of a method to extrapolate to new linkers and elaborations.

### Evaluation Metrics

We assessed the generated molecules with a range of 2D and 3D metrics, adopting a similar procedure to Imrie et al. ^17^. After first checking the generated molecules for validity, uniqueness, and novelty, we then determined if the generated examples were consistent with the 2D property filters used to produce the training set. While it is likely that there are many molecules that would fulfil the desired criteria of the user, the original molecule is a “true” correct answer and represents the best single ground truth available. As a result, a primary evaluation metric was the the recovery rate, which measures in how many cases the original molecule was recovered by the generation process.

Molecules which passed the 2D property filters were assessed on the basis of their 3D shape. We calculated 3D similarity by scoring conformers of the generated molecules against the original molecule using the same 3D shape and colour score utilised in Imrie et al. ^17^, based on the methods described in Putta et al. ^38^ and Landrum et al. ^39^. For both linker design and scaffold elaboration, we primarily assessed the 3D complementarity of the generated molecular component only (i.e. the linker or R-group) with the reference structure (SC_RDKit_ Generated). This score ranges between 0 (no match) and 1 (perfect match). Scores above 0.6 indicate a good match, while scores above 0.9 suggest an almost perfect match.

The focus of our analysis is based on SC_RDKit_ Generated as it directly captures the chemical differences between the proposed molecules. However, for linker design we also calculated the 3D metrics utilised in Imrie et al. ^17^ (SC_RDKit_ Molecule, SC_RDKit_ Fragments, RMSD) to ensure that the proposed linkers satisfy the basic structural constraints. We did not use these metrics in the scaffold elaboration experiments since the conformation of the molecular core is typically largely unaffected by its side chains.^40^

For each proposed compound, we generated 3D conformers using RDKit,^41^ adopting the filtering and sampling procedure proposed by Ebejer et al. ^42^. To calculate SC_RDKit_ Generated, we generated conformers in a constrained manner, biasing conformations towards those that maintained the conformation of the starting structure(s). However, to mitigate the risk of generating physically unrealistic structures, we removed high energy conformers. We then scored all conformers, taking the best score as the final score for a particular molecule.

### Comparison to Other Methods

For both linker design and scaffold elaboration, we compared DEVELOP to DeLinker^17^ and a version of the DeLinker method which is provided with the number of each pharmacophoric feature that should be present in the generated linker (“DeLinker-Counts”). The difference between DEVELOP and these two baselines is the structural information, ***D***, included in the feature vector, 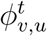. This comparison allowed us to assess directly the impact of (1) including pharmacophoric constraints, and (2) providing these constraints as a physically-meaningful 3D structural representation rather than a 1D count vector.

We also compared our results to recent deep learning methods for these design problems. For linker design, we compared our method to SyntaLinker,^18^ while for scaffold elaboration, we benchmarked against Arús-Pous et al. ^21^ (“REINVENT”). Both methods adopt a SMILES-based formulation and neither framework incorporates 3D information in the design process. In both cases, we retrained these models on the training sets described above using the open-source implementations provided by the authors to ensure a fair comparison between the methods tested. We adopted the same settings and hyperparameters described in the original publications.

### Experimental Setup

In all of our experiments, we used the same training sets (one for linker design and one for scaffold elaboration) derived from the ZINC data set to train all of the models considered. When evaluating using the data sets derived from CASF and PDBbind, we generated 250 molecules for each example for each of the methods considered. For the graph-based models (DeLinker, DeLinker-Counts, and DEVELOP), the number of atoms was set equal to the number of atoms in the original molecule. The pharmacophoric information provided to DeLinker-Counts and DEVELOP was derived directly from the ground truth molecule. In the case of SyntaLinker, the model was provided with the shortest linker bond distance.

## Results and Discussion

We validate the ability of our deep generative model (DEVELOP) to perform linker design and scaffold elaboration using 3D pharmacophoric information, reporting significant improvement over all other methods. Through the use of several canonical examples, we demonstrate the impact of the pharmacophoric constraints on the generated molecules. We show a significant improvement in the quality of generated molecules in large-scale evaluations on test sets derived from CASF and PDBbind, further demonstrating the importance of including pharmacophoric information. Finally, we illustrate the applicability of our approach to scaffold elaboration using an R-group optimisation case study derived from the literature.

### Importance of 3D Pharmacophoric Constraints

We assessed the impact of pharmacophoric constraints on the generation process empirically using two canonical examples for linker design and one example for scaffold elaboration (Table 1). The examples were all chosen from the PDBbind test sets (see Methods) and therefore none of the target elaborations were included in the training set. We generated 1000 molecules for each example using DeLinker, DeLinker-Counts, and DEVELOP.

**Table 1:**
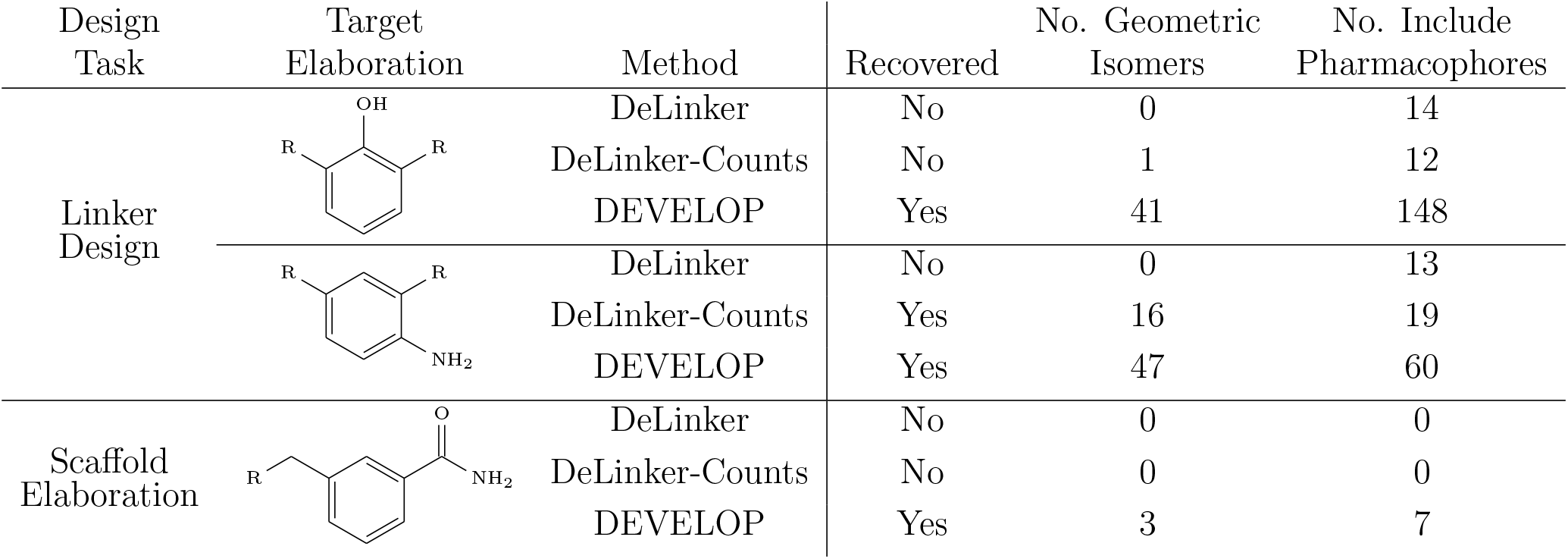
Impact of pharmacophoric constraints. Including 3D pharmacophoric information (DEVELOP) substantially improves the ability to generate molecules with desired interaction patterns.

Only DEVELOP was able to recover both of the canonical examples for linker design, with DeLinker not generating the correct linker in either case. The difference between the methods is further exemplified when considering geometric isomers with the same chemical structure but possibly different substitution patterns of the exit vectors and substituent. DEVELOP frequently generated linkers matching the chemical structure of the linker (41-47), while DeLinker did not produce a single geometric isomer.

The improved performance of DEVELOP is also evident when we assessed how many molecules included the desired pharmacophoric pattern of the examples (aromatic ring with correct substituent group). A significantly larger proportion of the generated molecules contained the desired pharmacophoric features when the 3D information was provided (60-148, DEVELOP) compared to not providing this information (13-14, DeLinker) or providing only 1D pharmacophore counts (12-19, DeLinker-Counts).

The largest difference in generated molecules occurred in the phenol example, where only one geometric isomer was generated by DeLinker and DeLinker-Counts combined compared to 41 from DEVELOP. This is a particularly difficult example for both DeLinker and DeLinker-Counts due to the presence of a donor-acceptor group, but illustrates the necessity of adopting a 3D representation.

The scaffold elaboration example proved challenging for all methods, primarily due to the size of the elaboration. Only DEVELOP recovered the 3-methyl-benzamide elaboration, with neither of the other two methods generating a single geometric isomer. In addition, DEVELOP was the only method to generate any elaborations containing the desired functionality of an aromatic system with an amide side-chain.

These examples demonstrate the importance of including pharmacophoric constraints for both linker design and scaffold elaboration. In all cases, it was only possible to consistently generate molecules with specific pharma-cophoric profiles when 3D pharmacophoric information was included.

### Linker Design Experiments On Large Test Sets

DEVELOP substantially outperformed all other methods on both the CASF and PDBbind test sets, with significant improvements in both the number of true linkers recovered and the proportion of generated molecules with high SC_RDKit_ Generated. This was achieved with limited impact on the uniqueness of the generated molecules and their ability to pass basic 2D chemical filters (Tables 2, S1, Figure 3).

**Table 2:** Linker design. PDBbind set results (see Methods, Evaluation Metrics for definitions of the metrics).

| Metric | SyntaLinker | DeLinker | DeLinker-Counts | DEVELOP |
| --- | --- | --- | --- | --- |
| Valid | 11.6% | 96.9% | 90.2% | 93.1% |
| Unique | 93.6% | 86.1% | 77.8% | 77.3% |
| Novel | 54.8% | 84.0% | 87.6% | 88.7% |
| Recovered | 0.3% | 1.9% | 8.7% | 22.4% |
| Pass 2D filters | 81.1% | 63.4% | 59.5% | 61.7% |
| SC<sub>RDKit</sub> Generated |  |  |  |  |
| >0.6 | 9.4% | 10.4% | 19.8% | 27.9% |
| >0.7 | 4.5% | 4.2% | 10.1% | 14.8% |
| >0.8 | 2.2% | 1.5% | 4.4% | 6.1% |
| >0.9 | 0.7% | 0.4% | 1.2% | 1.5% |

**Figure 3:**
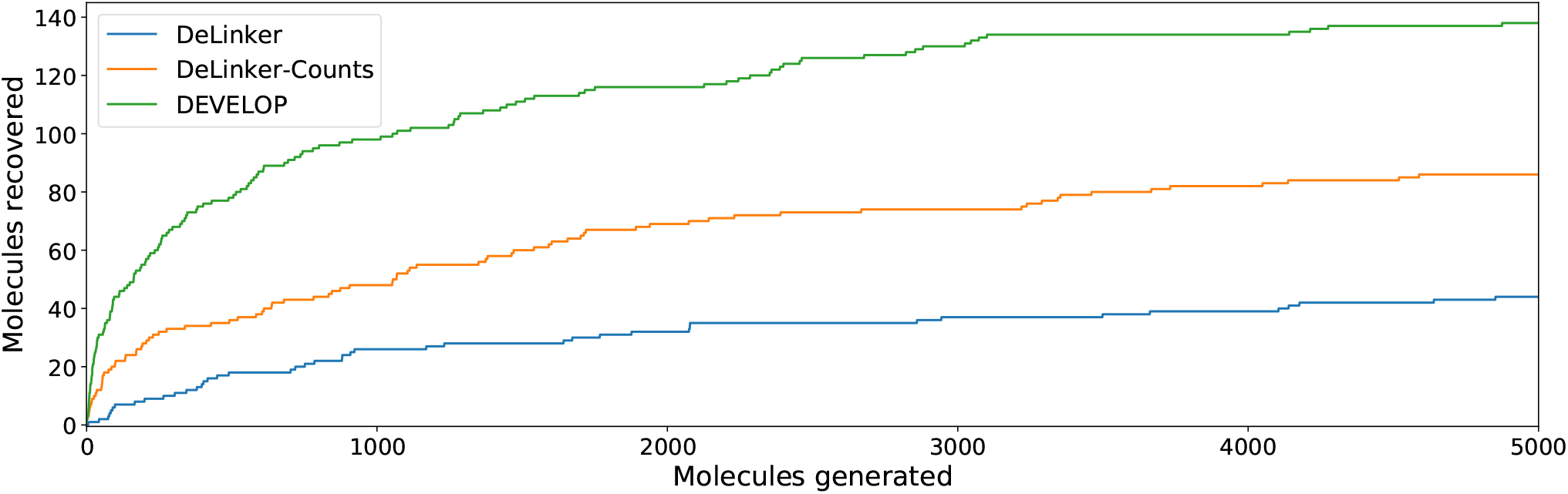
Linker design. Number of original molecules recovered as the number of generated molecules is increased. DEVELOP recovers the significantly more of the original molecules than both baselines for any number of samples generated.

In comparison, SyntaLinker performed poorly in particular as measured by the 2D metrics, producing weaker results than were reported in its original publication.^18^ SyntaLinker produced a low proportion of valid molecules and recovered only 0.3% of the original molecules. Due to the comparatively weak results, we focus the remainder of our analysis on the three graph-based methods. For further discussion on the SyntaLinker results, see the Supporting Information.

The proportion of valid molecules generated by the other three methods was high in all cases (>90%) with similar proportions of novel molecules proposed (69-71% on CASF, Table S1; 84-89% on PDBbind, Table 2). As is expected, as more structural information was provided to the model, the proportion of unique molecules decreased due to the constraints on the generative process. However, 58% and 77% of the molecules produced DEVELOP on CASF and PDBbind, respectively, were unique, demonstrating that the model still samples from chemical space and has not experienced mode collapse, degrading to a single or small number of solutions.

Incorporating pharmacophoric information substantially increased the recovery rate of the original molecules. On the CASF set, DEVELOP recovered 50% of the ground truth molecules, compared to 30% for DeLinker and 42% for DeLinker-Counts. The PDBbind set is particularly challenging with DeLinker only able to recover 1.9% of the original molecules, while a database-based method would not be able to recover any, due to there being no overlap with the training set. Including the count of each pharmacophore present in the original linker increased the proportion recovered to 8.7% (DeLinker-Counts). Crucially, providing this information as a 3D structural representation offered a significant benefit over simply providing the pharmacophore counts. On the PDBbind test set, DEVELOP recovered 22.4% of the original molecules, more than ten times as many as DeLinker and more than twice as many as DeLinker-Counts (Table 2).

A significant improvement is also seen when assessing the 3D similarity of the generated linkers to the original ones. DEVELOP improved the proportion of molecules with high structural similarity (SC_RDKit_ Generated > 0.8) by 300% and 39% compared to DeLinker and DeLinker-Counts, respectively, on the PDBbind test set (Table 2), with similar improvements on the CASF set (Table S1).

In addition to SC_RDKit_ Generated, we also calculated the 3D metrics employed in Imrie et al. ^17^, namely SC_RDKit_ Molecule, SC_RDKit_ Fragments, and RMSD. These metrics primarily capture whether the molecular linker allows the original substructures to adopt similar conformations, with the chemical features of the linker having limited to no effect on this score. The linkers generated by DEVELOP showed a substantial improvement on CASF compared to both DeLinker and DeLinker-Counts (Table S2), while performing similarly on PDBbind (Table S3).

As previously stated, these metrics primarily assess whether the linker can allow the starting substructures to adopt the required conformation. The additional information regarding the desired linker chemistry may well not improve these scores, even when linker quality is substantially improved.

To investigate whether the improvement in recovery rate is due to the number of linkers generated, we generated 5000 examples for each pair of starting fragments and assessed in how many cases the true linker was recovered (Figure 3). Due to sampling constraints, it was not possible to include SyntaLinker in this analysis (see the Supporting Information). The improvement in recovery rate of DEVELOP persisted even as substantially more linkers were generated. After several thousand examples, the rate of recovery of additional linkers decreased significantly for all methods, but remained the highest for DEVELOP. While increasing the number of samples further would be likely to yield more linkers being recovered, this effect may well be relatively small unless orders of magnitude more samples were generated. Figure 3 demonstrates that DEVELOP generates better linkers rather than simply producing similar molecules to DeLinker.

### Scaffold Elaboration Experiments On Large Test Sets

Large-scale assessments on the CASF and PDBbind test sets demonstrated that DEVELOP can effectively perform scaffold elaboration, with similar trends as the linker design experiments (Tables 3, S4 and Figure 4).

**Table 3:** Scaffold elaboration. PDBbind set results (see Methods, Evaluation Metrics for definitions of the metrics).

| Metric | REINVENT | DeLinker | DeLinker-Counts | DEVELOP |
| --- | --- | --- | --- | --- |
| Valid | 99.9% | 99.9% | 100.0% | 99.4% |
| Unique | 23.7% | 86.7% | 80.3% | 74.8% |
| Novel | 2.1% | 70.2% | 78.3% | 77.2% |
| Recovered | 0.0% | 1.4% | 4.4% | 14.9% |
| Pass 2D filters | 98.8% | 56.1% | 48.6% | 52.2% |
| SC<sub>RDKit</sub> Generated |  |  |  |  |
| >0.6 | 4.5% | 3.3% | 5.5% | 13.8% |
| >0.7 | 1.1% | 1.0% | 1.7% | 5.2% |
| >0.8 | 0.3% | 0.2% | 0.5% | 1.0% |
| >0.9 | 0.0% | 0.0% | 0.2% | 0.1% |

**Figure 4:**
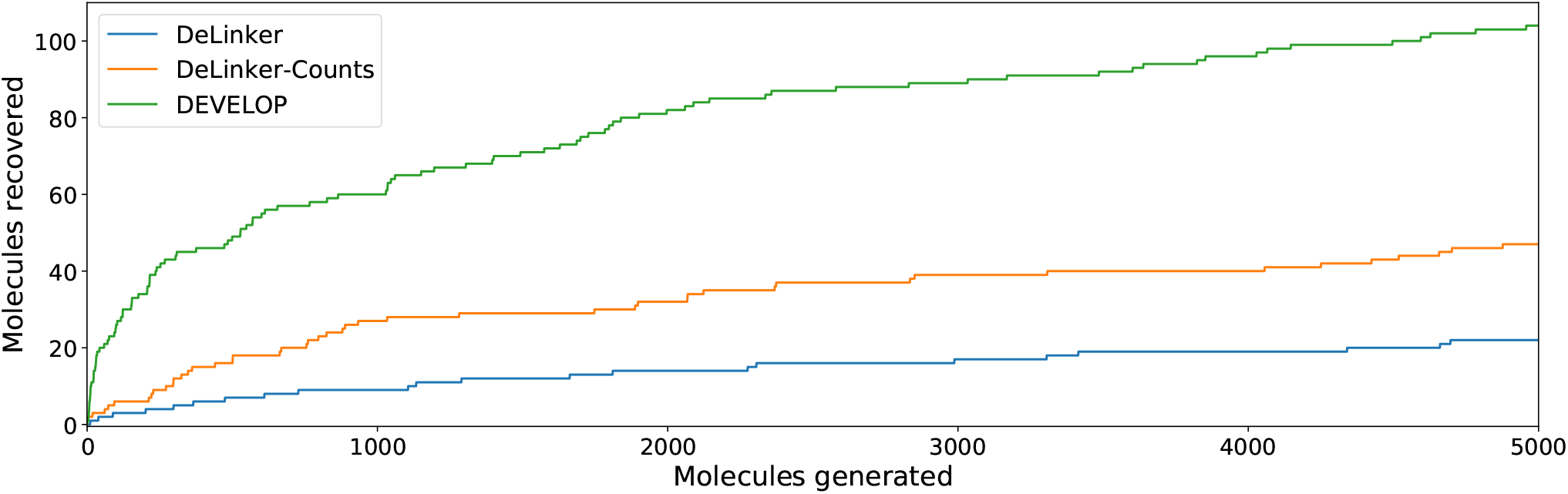
Scaffold elaboration. Number of original molecules recovered as the number of generated molecules is increased. DEVELOP recovers the significantly more of the original molecules than both baselines for any number of samples generated.

Almost all molecules generated by the graph-based models (DeLinker, DeLinker-Counts, and DEVELOP) are deemed valid since chemical valency is enforced during generation (the very small number of invalid molecules arises from cases where no elaboration is proposed, Table 3). The majority of molecules generated were unique, with uniqueness decreasing from 87% for DeLinker to 75% for DEVELOP as more constraints were provided. This was in line with expectations and mirrors the linker design experiments, with all methods proposing a high proportion of novel R-groups (35-43% on the CASF set, 70-78% on the PDBbind set).

In line with the performance for linker design, including 3D pharmacophoric information resulted in a substantially higher proportion of the true elaborations being recovered. On the CASF test set, DEVELOP recovered 69% of the ground truth molecules compared to 47% for DeLinker and 60% for DeLinker-Counts (Table S4). On the PDBbind set, DEVELOP recovered 15% of the original elaborations, an increase of ten-fold compared to DeLinker (1.4%, Table 3). This performance persisted as more molecules were generated (Figure 4). When 5000 elaborations were generated for each scaffold, DEVELOP recovered 35% of the original molecules compared to 16% when the 3D information was removed (DeLinker-Counts) and only 7% when no pharmacophoric information was included (DeLinker).

Finally, there was a substantial improvement in the 3D similarity of the generated molecules to the original ones. Of the elaborations which passed the 2D filters, 13.8% of those generated by DEVELOP obtained an SC_RDKit_ Generated score of greater than 0.6 compared to 3.3% and 5.5% obtained by DeLinker and DeLinker-Counts, respectively.

Almost none of the molecules generated by any method for the PDBbind test set achieved an SC_RDKit_ Generated score above 0.9. To reduce the impact of possible limitations of the conformer generation process, we recalculated SC_RDKit_ Generated using generated conformers of the ground truth molecules instead of the experimentally determined conformers (Tables S5, S6). On the PDBbind set, the proportion of generated molecules with SC_RDKit_ Generated > 0.9 remained low for all methods except DEVELOP, which increased to 2.6%. This represents a sizeable improvement over the next best method (DeLinker-Counts, 0.3%) and provides further validation of the improved quality of molecules generated by DEVELOP compared to the baselines.

REINVENT produced significantly fewer novel elaborations than either of the DeLinker models or DEVELOP, with only 2-3% of generated elaborations not contained in the training set (Table 3, S4). As such, REINVENT did not recover any of the original elaborations in the PDBbind test set, while on the CASF test set REINVENT recovered only 25% of the original elaborations compared to 69% for DEVELOP. In addition, REINVENT generated a similar proportion of elaborations that had high 3D similarity to the original molecules as DeLinker, and was substantially outperformed by both DeLinker-Counts and DEVELOP. This is expected given the additional information available to both models; however, it reinforces the importance of including pharmacophoric information.

### R-Group Optimisation Case Study

We further demonstrate the applicability of DEVELOP to R-group optimisation via a case study derived from the literature. Borkin et al. ^43^ developed a thienopyrimidine class of compounds to block the protein–protein interaction between menin and mixed lineage leukemia (MLL) fusion proteins. This interaction plays an important role in acute leukemias with MLL translocations, making this an important drug target. The authors’ previous work^44^ had led to the identification of a highly potent menin–MLL inhibitor (IC_50_=31 nM, GI_50_=0.55 *μ*M, PDB ID: 4X5Z) but required further improvement of cellular activity and drug-like properties to develop compounds with potential therapeutic value. This was achieved via structure-based optimisation of substituents introduced to the indole ring (Figure 5a).

**Figure 5:**
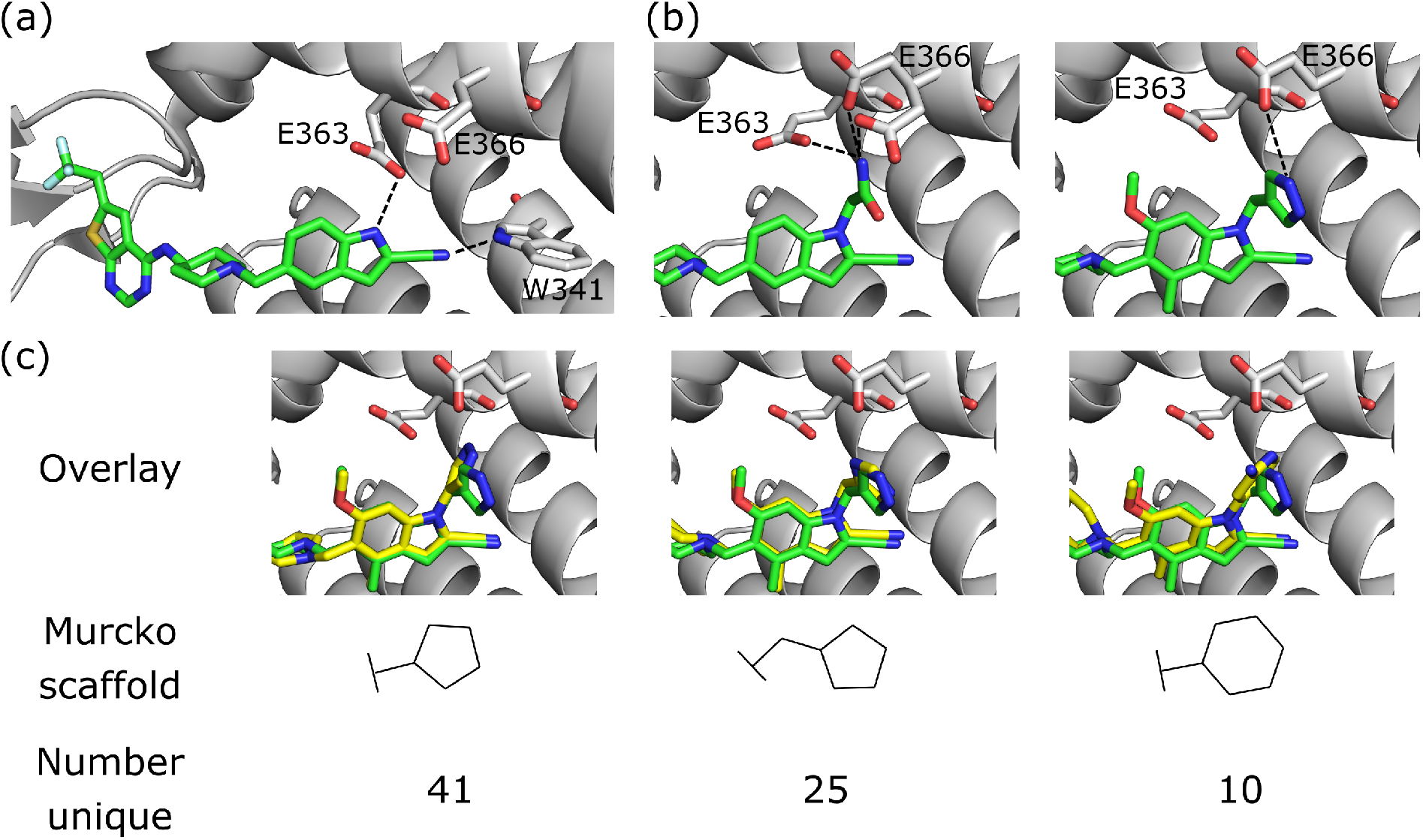
R-group optimisation case study. (a) Crystal structure (PDB ID 4X5Z) of the initial complex bound to menin. (b) Structure of two of the most potent optimised compounds (PDB IDs left 5DB2, right 5DB3). The dashed lines represent key interactions. (c) Overlay of the most potent optimised compound (green carbons, PDB ID 5DB3) and several compounds generated by DEVELOP (yellow carbons) that make similar hydrogen bonding interactions.

Following optimisation of several positions, the most potent compound displayed almost a seven-fold improvement in affinity in MLL-AF9 cells (GI_50_=83 nM, PDB ID: 5DB3, Figure 5b, right), while other highly potent compounds demonstrated favourable drug-like properties, such as significant improvements in selectivity, reduced lipophilicity, and bioavailability.

The most significant modification to the original compound was the optimisation of the hydrogen bond interactions with Glu363 and Glu366 on menin. The indole nitrogen in the original molecule was involved in a hydrogen bond with the side chain of Glu363 but was partially solvent exposed and was not forming interactions with Glu366 (Figure 5a). This led the authors to explore a variety of substituents containing hydrogen bond donors. Two potent substitutions were an acetamide group (Figure 5b, left) and 4-methylpyrazole (Figure 5b, right).

We investigated the ability of DEVELOP to propose R-groups that met the design hypothesis described in Borkin et al. ^43^. In particular, we sought to design both aromatic and non-aromatic hydrogen bond donor groups that were able to make similar interactions to the R-groups that were experimentally tested. We derived 3D pharmacophoric profiles from the ligands in PDB IDs 5DB2 and 5DB3 to serve as input to DEVELOP. For the pharmacophoric profile derived from 5DB2, we generated 1000 R-groups with a maximum of four, five, and six atoms, whilst for the pharma-cophoric profile derived from 5DB3 we generated 1000 molecules with a maximum of five, six, and seven atoms.

DEVELOP successfully recovered both of the experimentally-verified R-groups while generating many alternative molecules that could form similar interactions with menin. All methods were able to recover the acetamide R-group (Figure 5b, left). However, DEVELOP produced substantially more examples that matched the pharmacophoric profile (455) compared to both DeLinker-Counts (327) and DeLinker (103). All methods were also able to recover the 4-methylpyrazole R-group, although this elaboration was only generated once by DeLinker and DeLinker-Counts, compared to 61 times by DEVELOP. In addition, 237 of the elaborations generated by DEVELOP contained an aromatic system with a donor group linked to the indole via a methylene group compared to 50 for DeLinker and 11 for DeLinker-Counts.

We next sought to assess the alternatives to the pyrazole R-group (Figure 5b, right) that were proposed by DEVELOP. To validate the molecules proposed by DEVELOP, we docked the generated molecules containing an aromatic system and at least one donor group using GOLD^45^ and checked whether the docked pose formed hydrogen bonding interactions with Glu363 or Glu366. Three elaborations, together with their Murcko scaffolds, are shown in Figure 5c (yellow carbons) overlayed with the pyrazole R-group (green carbons). All of the examples appear to fit within the pocket and were able to form hydrogen bonds with Glu363 or Glu366, consistent with the stated design hypothesis.

## Conclusion

We have developed a method that combines GNNs with CNNs to incorporate 3D pharmacophoric constraints into molecular generation. Our approach allows prior knowledge to be used to control the design process and is readily extendable to alternate 3D structural representations.

We have demonstrated the applicability of our approach to both linker design and scaffold elaboration, two general tasks in the hit-to-lead and lead optimisation stages of drug discovery. The experimental results show that our model significantly outperforms previous methods for these problems and demonstrates the power of including pharmacophoric constraints as a 3D representation as opposed to a 1D count vector.

While the quality of the generated compounds has increased significantly, the problem of selecting which ones should be explored further remains a key consideration. Successful application of generative models relies on their successful integration into the broader drug discovery toolbox. An interesting development in this direction is described in Green et al. ^46^, who used CNNs to predict appropriate fragments given the structure of a protein-ligand complex. While their work was based on scoring a fixed database of fragments, extending such an approach to assess arbitrary elaborations could be readily combined with our method to rank generated molecules.

While the focus of our work is generating molecules with specific 3D characteristics, we do not directly assign atomic coordinates during generation. The direct generation of 3D molecular structures is an exciting development,^47,48^ but has not yet been applied to druglike molecules nor are existing methods directly applicable to the settings considered in this work. Extending our framework to generate atomic coordinates directly is an avenue for future work. Similarly, while we have shown encoding graph- and voxel-based representations separately is effective, unifying both with a single encoder that is 3D-aware could provide further benefit.

Finally, the pharmacophoric profiles used for our experiments were extracted from known molecules. While existing molecules can often be used as the basis for specifying desired pharmacophoric profiles in scaffold hopping or R-group optimisation, for fragment linking or elaboration a suitable ligand might not be available to derive a pharmacophoric profile, necessitating the manual specification of pharmacophoric features by a human expert. Accurate prediction of useful pharmacophoric features, directly from the protein structure or by other means, is therefore an important next step.

We believe that our method will allow greater synergy between human design hypotheses and machine learning-based molecular design. Code is available at https://github.com/oxpig/DEVELOP.

## Acknowledgement

F.I. is supported by funding from the Engineering and Physical Sciences Research Council (EPSRC) and Exscientia (Reference: EP/N509711/1). T.E.H. is supported by funding from EPSRC, LifeArc, F. Hoffmann-La Roche AG, and UCB Pharma (Reference: EP/L016044/1).

