## supplementary material for "Deep Generative Design with 3D Pharmacophoric Constraints"

### Supporting Information
### Deep Generative Design with 3D Pharmacophoric Constraints

Fergus Imrie,[†] Thomas E. Hadfield,[†] Anthony R. Bradley,[‡] and Charlotte M. Deane[*,†]

†*Oxford Protein Informatics Group, Department of Statistics, University of Oxford, Oxford OX1 3LB, UK*

‡*Exscientia Ltd, The Schrödinger Building, Oxford Science Park, Oxford OX4 4GE, UK*

### Additional DEVELOP model details

#### Atom types.

In line with both Liu et al.[1] and Imrie et al.[2], 14 atom types are permitted: carbon, nitrogen ($N^-$, N, $N^+$), oxygen ($O^-$, O, $O^+$), fluorine, chlorine, bromine, iodine, and sulphur (maximum valence 2, 4, or 6).

#### Hyperparameters.

As discussed in Methods, we used the same hyperparameters to train DEVELOP as adopted in Imrie et al.[2]. In particular, we trained the model with a learning rate of 0.001 for 10 epochs using the Adam optimiser and a batch size of 16. The hidden state dimension was set at 32, the encoding dimension 4, and $\lambda_{KL}$ 0.3. The same hyperparameters were used for all experiments.

### Comparison to SyntaLinker

SyntaLinker[3] is a transformer-based model for linker design that utilises a SMILES-based representation. The results in Table S1 are not directly comparable with the original publication since SyntaLinker was both trained and evaluated on different data sets in Yang et al.[3]. As described in Methods, we ensured a fair comparison between all methods by training and assessing all methods on the same data sets.

SyntaLinker produced substantially weaker results than were reported in its original publication.[3] In particular, SyntaLinker produced a very low proportion of valid molecules on both the CASF (Table S1) and PDBbind (Table 2) test sets. The poor validity of generated molecules can be explained by the sampling method used by SyntaLinker, which employs beam search[4] to generate SMILES strings. When sampling only the most likely sequences, the validity of generated molecules is relatively high; when the number of sequences sampled is increased, the validity falls significantly as lower probability sequences are selected. In addition, we note that this sampling procedure limits the number of molecules that can be generated by SyntaLinker. However, as a result of using beam search, almost all of the SMILES strings generated by SyntaLinker correspond to unique molecules.

In addition to the low proportion of valid molecules generated, SyntaLinker recovered only 8% of the original molecules on the CASF set compared to 30% for DeLinker and 50% for DEVELOP, while on the PDBbind set SyntaLinker recovered 0.3% of the original molecules compared to 1.9% and 22.4% for DeLinker and DEVELOP, respectively. The 3D shape similarity of the molecules generated by SyntaLinker was significantly lower than DEVELOP for both the CASF (Table S1) and PDBbind (Table 2) test sets.

### Additional results

Table S1: Linker design. CASF set results.

| Metric | SyntaLinker | DeLinker | DeLinker-Counts | DEVELOP |
|---|---|---|---|---|
| Valid | 9.8% | 94.7% | 86.0% | 89.6% |
| Unique | 95.1% | 72.9% | 58.6% | 58.2% |
| Novel | 54.4% | 68.7% | 68.4% | 71.1% |
| Recovered | 7.5% | 29.8% | 41.5% | 50.0% |
| Pass 2D filters | 80.5% | 71.7% | 71.7% | 68.6% |
| $SC_{RDKit}$ Generated | | | | |
| >0.6 | 14.0% | 23.0% | 40.6% | 45.5% |
| >0.7 | 7.0% | 12.2% | 27.9% | 31.0% |
| >0.8 | 3.8% | 6.4% | 18.4% | 20.7% |
| >0.9 | 1.5% | 2.7% | 10.3% | 12.2% |

Table S2: Linker design. Alternative 3D similarity metrics for CASF test set.

| Metric | SyntaLinker | DeLinker | DeLinker-Counts | DEVELOP |
|---|---|---|---|---|
| $SC_{\mathrm{RDKit}}$ Molecule | | | | |
| >0.7 | 22.5% | 16.3% | 22.5% | 25.5% |
| >0.8 | 4.7% | 3.6% | 7.0% | 7.2% |
| >0.9 | 0.5% | 0.8% | 2.9% | 3.0% |
| $SC_{\mathrm{RDKit}}$ Fragments | | | | |
| >0.7 | 42.6% | 38.7% | 40.8% | 43.0% |
| >0.8 | 14.2% | 12.3% | 14.7% | 15.4% |
| >0.9 | 1.3% | 1.6% | 3.2% | 3.8% |
| RMSD | | | | |
| <1.00Å | 30.5% | 26.6% | 28.1% | 30.7% |
| <0.75Å | 11.8% | 9.3% | 10.6% | 12.9% |
| <0.50Å | 3.0% | 2.4% | 4.3% | 4.7% |

Table S3: Linker Design. Alternative 3D similarity metrics for PDBbind test set.

| Metric | SyntaLinker | DeLinker | DeLinker-Counts | DEVELOP |
|---|---|---|---|---|
| $SC_{\mathrm{RDKit}}$ Molecule | | | | |
| >0.7 | 12.6% | 12.8% | 15.8% | 17.3% |
| >0.8 | 2.4% | 1.9% | 3.3% | 3.7% |
| >0.9 | 0.1% | 0.0% | 0.2% | 0.2% |
| $SC_{\mathrm{RDKit}}$ Fragments | | | | |
| >0.7 | 35.0% | 34.5% | 35.7% | 34.0% |
| >0.8 | 11.8% | 9.9% | 10.2% | 10.3% |
| >0.9 | 2.0% | 1.1% | 0.8% | 1.0% |
| RMSD | | | | |
| <1.00Å | 26.7% | 24.8% | 26.1% | 25.0% |
| <0.75Å | 10.3% | 8.5% | 8.9% | 8.7% |
| <0.50Å | 3.0% | 1.4% | 1.4% | 1.6% |

Table S4: Scaffold elaboration. CASF set results.

| Metric | REINVENT | DeLinker | DeLinker-Counts | DEVELOP |
|---|---|---|---|---|
| Valid | 99.9% | 99.9% | 99.8% | 99.1% |
| Unique | 27.4% | 56.6% | 39.1% | 29.1% |
| Novel | 3.2% | 43.4% | 40.9% | 34.5% |
| Recovered | 25.3% | 47.3% | 59.9% | 68.8% |
| Pass 2D filters | 98.2% | 72.6% | 75.1% | 80.0% |
| $SC_{RDKit}$ Generated | | | | |
| >0.6 | 16.0% | 13.7% | 27.6% | 43.9% |
| >0.7 | 9.3% | 7.6% | 19.6% | 32.3% |
| >0.8 | 5.2% | 4.0% | 12.9% | 20.9% |
| >0.9 | 1.8% | 2.0% | 6.0% | 9.4% |

Table S5: Scaffold elaboration. CASF set results, $SC_{RDKit}$ Generated calculated with RDKit generated reference conformers.

| Metric | REINVENT | DeLinker | DeLinker-Counts | DEVELOP |
|---|---|---|---|---|
| Valid | 99.9% | 99.9% | 99.8% | 99.1% |
| Unique | 27.4% | 56.6% | 39.1% | 29.1% |
| Novel | 3.2% | 43.4% | 40.9% | 34.5% |
| Recovered | 25.3% | 47.3% | 59.9% | 68.8% |
| Pass 2D filters | 98.2% | 72.6% | 75.1% | 80.0% |
| $SC_{RDKit}$ Generated | | | | |
| >0.6 | 25.0% | 21.0% | 37.5% | 58.0% |
| >0.7 | 17.6% | 14.0% | 29.8% | 48.8% |
| >0.8 | 10.5% | 7.5% | 21.5% | 36.6% |
| >0.9 | 4.7% | 4.5% | 15.0% | 26.9% |

Table S6: Scaffold elaboration. PDBbind set results, $SC_{RDKit}$ Generated calculated with RDKit generated reference conformers.

| Metric | REINVENT | DeLinker | DeLinker-Counts | DEVELOP |
|---|---|---|---|---|
| Valid | 99.9% | 99.9% | 100.0% | 99.4% |
| Unique | 23.7% | 86.7% | 80.3% | 74.8% |
| Novel | 2.1% | 70.2% | 78.3% | 77.2% |
| Recovered | 0.0% | 1.4% | 4.4% | 14.9% |
| Pass 2D filters | 98.8% | 56.1% | 48.6% | 52.2% |
| $SC_{RDKit}$ Generated | | | | |
| >0.6 | 10.5% | 8.4% | 14.1% | 29.7% |
| >0.7 | 4.4% | 3.1% | 6.1% | 14.3% |
| >0.8 | 0.8% | 0.9% | 2.1% | 6.9% |
| >0.9 | 0.1% | 0.1% | 0.3% | 2.6% |
